# Ensuring that fundamentals of quantitative microbiology are reflected in microbial diversity analyses based on next-generation sequencing

**DOI:** 10.1101/2021.06.19.449110

**Authors:** Philip J. Schmidt, Ellen S. Cameron, Kirsten M. Müller, Monica B. Emelko

## Abstract

Diversity analysis of amplicon sequencing data is mainly limited to plug-in estimates calculated using normalized data to obtain a single value of an alpha diversity metric or a single point on a beta diversity ordination plot for each sample. As recognized for count data generated using classical microbiological methods, read counts obtained from a sample are random data linked to source properties by a probabilistic process. Thus, diversity analysis has focused on diversity of (normalized) samples rather than probabilistic inference about source diversity. This study applies fundamentals of statistical analysis for quantitative microbiology (e.g., microscopy, plating, most probable number methods) to sample collection and processing procedures of amplicon sequencing methods to facilitate inference reflecting the probabilistic nature of such data and evaluation of uncertainty in diversity metrics. Types of random error are described and clustering of microorganisms in the source, differential analytical recovery during sample processing, and amplification are found to invalidate a multinomial relative abundance model. The zeros often abounding in amplicon sequencing data and their implications are addressed, and Bayesian analysis is applied to estimate the source Shannon index given unnormalized data (both simulated and real). Inference about source diversity is found to require knowledge of the exact number of unique variants in the source, which is practically unknowable due to library size limitations and the inability to differentiate zeros corresponding to variants that are actually absent in the source from zeros corresponding to variants that were merely not detected. Given these problems with estimation of diversity in the source even when the basic multinomial model is valid, sample-level diversity analysis approaches are discussed.

**Highlights:**

- Random error in amplicon sequencing method should be considered in diversity analysis
- Clustering, amplification, and differential recovery distort sample diversity
- The multinomial model for compositional count data is compromised by amplification
- There are three types of zeros in amplicon sequencing data, including missing zeros
- Source alpha diversity estimates are biased by unknown number of unique variants

## 1.0 Introduction

Analysis of microbiological data using probabilistic methods has a rich history, with examination of both microscopic and culture-based data considered by prominent statisticians a century ago (e.g., Student, 1907; Fisher et al., 1922). The most probable number method for estimating concentrations from suites of presence-absence data is inherently probabilistic (e.g., McCrady, 1915), though routine use of tables (or more recently software) obviates consideration of the probabilistic link between raw data and the estimated values of practical interest. Both the analysis of microbiological data and the control of the methods through which such data are obtained are grounded in statistical theory (e.g., Eisenhart & Wilson, 1943). More recently, the issue of estimating microbial concentrations and quantifying the uncertainty therein when some portion of microorganisms gathered in an environmental sample are not observed by the analyst has added to the complexity of analyzing microscopic enumeration data (e.g., Emelko et al., 2010). These examples share the common theme that the concentration of microorganisms in some source of interest is indirectly and imprecisely estimated from the discrete data produced by microbiological examination of samples (e.g., counts of cells/colonies or the number of aliquots exhibiting bacterial growth). The burgeoning microbiological analyses grounded in polymerase chain reactions (Huggett et al., 2015) likewise feature discrete objects (specific sequences of genetic material) that are prone to losses in sample processing, but these methods are further complicated by the variability introduced through amplification and reading (e.g., fluorescence signals or sequencing).

In next-generation amplicon sequencing, obtained data consist of a large library of nucleic acid sequences extracted and amplified from environmental samples, which are then tabulated into a set of counts associated with amplicon sequence variants (ASVs) or some grouping thereof (Callahan et al., 2017). The resulting data are regarded as a quantitative representation of the relative abundance (i.e., proportions) of various organisms in the source rather than absolute abundance (i.e., concentrations), thus leading to compositional data (Gloor et al., 2017). Among the many categories of analyses performed on such data are (1) differential abundance analysis to compare proportions of particular variants among samples and their relation to possible covariates and (2) diversity analysis that concerns the number of unique variants detected, how the numbers of reads vary among them, and how these characteristics vary among samples (Calle, 2019). Conventional analysis of these data is confronted with several problems (McMurdie & Holmes, 2014; Kaul et al., 2017; McKnight et al., 2018): (1) a series of samples can have diverse library sizes (i.e., numbers of sequence reads), motivating “normalization”, (2) there are many normalization approaches from which to choose, and (3) many normalization and data analysis approaches are complicated by large numbers of zeros in ASV tables. These issues can be overcome in differential abundance analysis through use of probabilistic approaches such as generalized linear models (e.g., McMurdie and Holmes, 2014) that link raw ASV count data and corresponding library sizes to a linear model without the need for normalization or special treatment of zeros. Diversity analysis, however, is more complicated because the amount of diversity exhibited in a particular sample (alpha diversity) or apparent similarity or dissimilarity among samples (beta diversity) is a function of library size (Hughes and Hellmann, 2005), and methods to account for this are not standardized.

A variety of methods have been applied to prepare amplicon sequencing data for downstream diversity analyses, most of which involve some form of normalization. Normalization options include (1) rarefying that randomly subsamples from the observed sequences to reduce the library size of a sample to some normalized library size shared by all samples in the analysis (Sanders, 1968), (2) simple proportions (McKnight et al., 2019), and (3) a continually expanding set of data transformations such as centered-log ratios (e.g., Gloor et al., 2017), geometric means pairwise ratio (e.g., Chen et al., 2018) or variance stabilizing transformations (e.g., Love et al., 2014). Rarefying predates high throughput sequencing methods (including applications beyond sequencing of the 16S rRNA gene such as RNA sequencing) and originated in traditional ecology. Statistically, these approaches to estimation of sample diversity in the source treat manipulated sample data as a population because the non-probabilistic analysis of a sample (called a plug-in estimate) leads to a single diversity value or a single point on an ordination plot.

While it would increase computational complexity to do so, it is more theoretically sound to acknowledge that the observed library of sequence reads in a sample is an imperfect representation of the diversity of the source from which the sample was collected and that no one-size-fits-all normalization of the data can remedy this. ASV counts would then be regarded as a suite of random variables that are collectively dependent on the sampling depth (library size) and underlying simplex of proportions that can only be imperfectly estimated from the available data. Analysis of election polls is somewhat analogous in that it concerns inference about the relative composition (rather than absolute abundance) of eligible voters who prefer various candidates. A key distinction is that such analysis does not presume that the fraction of respondents favouring a particular candidate or party (or some numerical transformation thereof) is an exact measurement of the composition of the electorate. Habitual reporting of a margin of error with proportional results (Freedman et al., 1998) exemplifies that such polls are acknowledged to be samples from a population in which the small number of eligible voters surveyed is central to interpretation of the data. Willis (2019) applies this approach to thinking about amplicon sequencing data in the estimation of alpha diversity by estimating diversity in a source from sample data using knowledge about random error to characterize uncertainty in source diversity.

Here, (1) the random process yielding amplicon sequencing data believed to be representative of microbial community composition in the source and (2) how this theory contributes to estimating the Shannon index alpha diversity metric using such data, particularly when library sizes differ and zero counts abound, are examined in detail. Theory applied to estimate microbial concentrations in water from data obtained using classical microbiological methods is extended to this type of microbiological assay to describe both the types of error that must be considered and a series of mechanistic assumptions that lead to a simple statistical model. The mechanisms leading to zeros in amplicon sequencing data and common issues with how zeros are analyzed in all areas of microbiology are discussed. Bayesian analysis is evaluated as an approach to drawing inference from a sample library about alpha diversity in the source with particular attention to the meaning and handling of zeros. This work addresses a path to evaluating microbial community diversity given the inherent randomness of amplicon sequencing data. It is based on established fundamentals of quantitative microbiology and provides a starting point for further investigation and development.

## 2.0 Describing and modelling errors in amplicon sequencing data

A theoretical model for the error structure in microbial data can be developed by contemplating the series of mechanisms introducing variability to the number of a particular type of microorganism (or gene sequence) that are present in a sample and eventually observed. This prioritizes understanding how random data are generated from the population of interest (e.g., the source microbiome) over the often more immediate problem of how to analyze a particular set of available data. Probabilistic modelling is central to such approaches, not just a data analytics tool. Rather than reviewing and attempting to synthesize the various probabilistic methods that have been applied to amplicon sequencing, the approach herein builds on a foundation of knowledge surrounding random errors in microscopic enumeration of waterborne pathogens (e.g., Nahrstedt & Gimbel, 1996; Emelko et al., 2010) to address the inherently more complicated errors in amplicon sequencing data. This study addresses the foundational matter of inferring a source microbiome alpha diversity metric from an individual sample because dealing with more complex situations inherent to microbiome analysis requires a firm grasp of such simple scenarios. Accordingly, hierarchical models for alpha diversity analyses that link samples to a hypothetical meta-community (e.g., McGregor et al., 2020) and approaches for differential abundance analysis in which the covariation of counts of several variants among multiple samples may be a concern (e.g., Mandal et al., 2015) are beyond the scope of this work. When random errors in the process linking observed data to the population characteristics of interest are integrated into a probabilistic model, it is possible to apply the model in a forward direction to simulate data given known parameter values or in a reverse direction to estimate model parameters given observed data. This reversibility is harnessed later in this paper to simulate data from a hypothetical source and evaluate how well Bayesian analysis of those data estimates the actual Shannon index of the source.

### 2.1 Describing amplicon sequencing data as a random sample from an environmental source

Microbial community analysis involves the collection of samples from a source such as environmental waters or the human gut (Shokralla et al., 2012). This study addresses the context of water samples because the plausibility that some sources could be homogeneous provides a comparatively simple and well understood statistical starting point for modelling—many other microbiomes are inherently not well mixed. When a sample is collected, it is presumed to be representative of some spatiotemporal portion of a water source such as a particular geographic location and depth in a water body and time of sampling. A degree of local homogeneity surrounding the location and time of the collected sample is often presumed so that randomness in the number of a particular type of microorganism contained in the sample (random sampling error) would be Poisson-distributed with mean equal to the product of concentration and volume. There are many reasons for which a series of samples presumed to be replicates from a particular source may yield microorganism counts that are over-dispersed relative to such a Poisson distribution (Schmidt et al., 2014), including (1) clustering of microorganisms to each other or on suspended particles, (2) spatiotemporally variable concentration, (3) variable volume analyzed, and (4) errors in sample processing and counting of microorganisms. Variable concentration and inconsistent sample volumes are not considered herein because the focus is on relative abundance (i.e., not estimation of concentrations) and samples that are not presumed to be replicates (i.e., analysis focuses on individual samples). Non-random dispersion could be a concern affecting estimates of diversity and relative abundance because clustering may inflate variability in the counts of a particular microorganism. For example, clustering could polarize results between unusually large numbers if a large cluster is captured and absence otherwise rather than yielding a number that varies minimally around the average.

The remainder of this analysis focuses on errors in sample handling and processing, nucleic acid amplification, and gene sequence counting. To be representative of relative abundance of microorganisms in the source, it is presumed that a sample is handled so that the community in the analyzed sample is compositionally equivalent to the community in the sample when it was collected (Fricker et al., 2019). Any differential growth or decay among types of microorganisms will bias diversity analysis. A series of sample processing steps is then needed to extract and purify the nucleic material so that the sample is reduced to a size and condition ready for PCR (polymerase chain reactions). Losses may occur throughout this process, such as adhesion to glassware, residuals not transferred, failure to extract nucleic material from cells, and sample partitioning during concentration and/or purification steps. These introduce random analytical error (because a method with 50% analytical recovery cannot recover 50% of one discrete microorganism, for example), and likely also non-constant analytical recovery if the capacity of the method to recover a particular type of microorganism varies randomly from sample to sample (e.g., 60% in one sample and 40% in the next). Any differential analytical recovery among types of microorganisms (e.g., if one type of microorganism is more likely to be successfully observed than another) will bias diversity analysis of the source. Varying copy numbers of genes among types of microorganisms as well as genes associated with non-viable organisms can also bias diversity analysis. PCR amplification is then performed with specific primers to amplify targeted genes, which may not perfectly double the number of gene copies in each cycle due to various factors including primer match. Any differential amplification efficiency among types of microorganisms will bias diversity analysis of the source, as will amplification errors that produce and amplify variants that do not exist in the source (unless these are readily identified and removed from sequencing data). Finally, the generated library of sequence reads is only a subsample of the sequences present in the amplified sample. Production of sequences that are not present in the original sample (e.g., chimeric sequences, misreads) is a form of loss if they detract from sequences that ought to have been read instead, and the resulting sequences may not be perfectly removed from the data (either failing to remove invalid sequences or erroneously removing valid sequences). Any differential losses at this stage will once again bias diversity analysis of the source, as will inadvertent inclusion of false sequences. Thus, the number of microorganisms gathered in a sample, the number of genes successfully reaching amplification, the number of genes after amplification, and the number of genes successfully sequenced are all random. Due to this collection of unavoidable random errors, the validity of diversity analysis approaches that regard samples (or normalized transformations of them) as exact compositional representations of the source requires further examination.

### 2.2 Modelling random error in amplicon sequencing data

For all of the reasons described above, it is impractical to regard libraries of sequence reads as indicative of absolute abundance in the source. We suggest that it is also impossible to regard them as indicative of relative abundance in the source without acknowledging a suite of assumptions and carefully considering what effect departure from those assumptions might have. By presuming that sequence reads are generated independently based on proportions identical to the proportional occurrence of those sequences in the source from which the sample was collected, the randomness in the set of sequence reads will yield a multinomial distribution. [For large random samples from small populations, a multivariate hypergeometric model without replacement may be more appropriate]. This is analogous to election poll data (if the poll surveys a small random sample of voters from a large electorate), repeatedly rolling a die, or repeatedly drawing random lettered tiles from a bag with replacement. The binary equivalent is a binomial model, which may form the basis of logistic regression to describe the proportion of sequences of a particular type as a function of possible covariates, recognizing how count data are random variables depending on respective library sizes and underlying proportions of interest.

Multinomial models are foundational to probabilistic analysis of count-based compositional data (e.g., McGregor et al., 2020), but mechanisms through which natural variability arises in the source (such as microorganism dispersion) and the sample collection and processing methodology (such as losses, amplification, and subsampling) must be considered because they may invalidate such a model for amplicon sequencing data—these need to be considered. Table 1 summarizes the random errors discussed above, contextualizes them in terms of compatibility with the multinomial relative abundance model, and summarizes the assumptions that must be made to use a multinomial model.

**Table 1:** Summary of random errors in amplicon sequencing and associated assumptions in the multinomial relative abundance model

| Error source | Description of error and compatibility with multinomial model | Assumptions |
| --- | --- | --- |
| Sample collection | The random sampling error describing variability in the number of discrete objects captured in a sample yields a Poisson distribution if microorganisms are randomly dispersed in a large source. This error is compatible with a multinomial model for proportional abundance of variants. Clustering, including multiple gene copies per organism, leads to excess variability that is incompatible with a multinomial model. | • All microorganisms are randomly dispersed (i.e., not clustered) with only one gene copy each* |
| Sample handling | The number of a particular type of microorganism may increase or decrease between sample collection and sample processing. Growth inflates the number of microorganisms at the level of diversity represented before growth occurred and is incompatible with a multinomial model. Decay is a form of random analytical error that is compatible with a multinomial model if it is consistent among variants. | • No growth<br>• No differential decay (analytical recovery) among variants |
| Sample processing | The number of gene sequences subjected to amplification may be lower than the number in the sample prior to processing due to losses (e.g., adherence to apparatus, not all genes extracted, sample partitioning). This is compatible with a multinomial model if analytical recovery is constant among variants. | • No differential losses (analytical recovery) among variants |
| Amplification | The number of gene sequences is purposefully increased using polymerase chain reactions, inflating the number of gene sequences at the level of diversity represented before amplification occurred, and is incompatible with a multinomial model. Copy errors are a form of loss for the original sequences that were incorrectly copied and produces erroneous sequences that may then be further amplified. Erroneous sequences are incompatible with a multinomial model unless all of them are removed from the data. | • Pre-amplification variant diversity is fully identical to source diversity and sequences are perfectly duplicated in each PCR cycle*<br>• No differential amplification efficiency or potential for copy errors among variants<br>• Data denoising must remove all erroneous sequence reads and no legitimate reads |
| Amplicon sequencing | Only a subsample of sequences are read, and all variants must be equally likely to be read. Sequence reading errors are a form of loss for the original sequences that were incorrectly read and also produces erroneous sequence reads. Sequence reading errors are incompatible with a multinomial model unless all resultant erroneous sequences are removed from the data. | • No differential sequence reading errors among variants or differential losses<br>• Data denoising must remove all erroneous sequence reads and no legitimate reads |

211 \*Without these difficult assumptions, the multinomial model describes post-amplification variant diversity rather than source microbial diversity

Based on some simulations (see R code in Supplementary Material), it was determined that random sampling error consistent with a Poisson model is compatible with the multinomial relative abundance model (using the binomial model as a two-variant special case). Specifically, this featured Poisson-distributed counts of two variants with means following a 2:1 ratio and graphical evidence that this process is consistent with a binomial model (also with a 2:1 ratio of the two variants) when the result was conditioned on a particular library size. It must be noted that this is not a formal proof, as “proof by example” is a logical fallacy (unlike “disproof by counter-example”). Critically, clustering of gene copies in the source causes the randomness in sequence counts to depart from a multinomial model, as proven by simulation in the Supplementary Material (following a disproof by counter-example approach). When the above process was repeated with counts following a negative binomial model that is over-dispersed with respect to the Poisson model, the variation in counts conditional on a particular library size was no longer consistent with the binomial model. Microorganisms having multiple gene copies is a form of clustering that invalidates the model.

Any form of loss or subsampling is compatible with the multinomial model so long as it affects all sequence variants equally. If each of a set of original proportions is multiplied by the same weight (analytical recovery), then the set of proportions adjusted by this weighting is identical to the original proportions (e.g., a 2:1 ratio is equal to a 1:0.5 ratio if all variants have 50% analytical recovery).

Growth and amplification must also not involve differential error among variants, but even in absence of differential error they have an important effect on the data and evaluation of microbiome diversity. These processes inflate the number of sequences present, but only with the potentially reduced or atypical diversity represented in the sample before such inflation. For example, a hypothetical sample with 100 variants amplified to 1000 will have the diversity of a 100-variant sample in 1000 reads, which may inherently be less than the diversity of a 1000-variant sample directly from the source. Amplification fabricates additional data in a process roughly opposite to discarding sequences in rarefaction; it resamples from a small pool of genetic material to make more whereas rarefaction subsamples from a larger pool of gene sequences to yield less (i.e., a smaller library size). Based on some simulations (see R code in Supplementary Material), it was proven that amplification is incompatible with the multinomial relative abundance model (following a disproof by counter-example approach). Specifically, the distribution of counts when two variants with a 2:1 ratio are amplified from a library size of four to a library size of six, the results differ from the distribution of counts obtained from a binomial model.

Representativeness of source diversity and compatibility with the multinomial relative abundance model can only be assured if the post-amplification diversity happens to be fully identical to the pre-amplification diversity and the observed library is a small random sample of the amplified genetic material. Such an assumption may presume random happenstance more so than a plausible probabilistic process, though it would be valid in the extreme special case where pre-amplification diversity is fully identical to source diversity and every sequence is perfectly duplicated in each cycle (with no erroneous sequences produced). Without making relatively implausible assumptions or having detailed understanding and modelling of the random error in amplification, observed libraries are only representative of post-amplification diversity and indirectly representative of source diversity. This calls into question the theoretical validity of multinomial models as a starting point for inference about the proportional composition of microbial communities using amplification-based data. Nonetheless, the multinomial model was used as part of this study in some illustrative simulation-based diversity analysis experiments.

## 3.0 The many zeros of amplicon sequencing data

As in other fields (Helsel, 2010), zeros in microbiology have led to much ado about nothing (Chik et al., 2018). They are (1) commonly regarded with skepticism that is hypocritical of non-zero counts (e.g., assuming that counts of zero result from error while counts of two are infallible), (2) often substituted with non-zero values or omitted from analysis altogether, and (3) a continued subject of statistical debate and special attention (such as detection limits and allegedly censored microbial data). Careful consideration of zeros is particularly relevant to diversity analysis of amplicon sequencing data because they often constitute a large portion of ASV tables. They may or may not appear in sample-specific ASV data, but they often appear when the ASV table of several samples is filled out (e.g., when an ASV that appears in some samples does not appear in others, zeros are assigned to that ASV in all samples in which it was not observed). They may also be created by zeroing singleton reads (Callahan et al., 2016), but this issue (and the bias arising if some singletons are legitimate read counts) is not specifically addressed in this study. Zeros often receive special treatment during the normalization step of compositional microbiome analysis (Thorsen et al., 2016; Tsilimigras and Fodor, 2016; Kaul et al., 2017), including removal of rows of zeros and fabrication of pseudo-counts with which zeros are substituted (to enable logarithmic transformations among other reasons).

We propose a classification of three types of zeros: (1) non-detected sequences (also caused sampling zeros), (2) truly absent sequences (also called structural zeros), and (3) missing zeros. This differs from the three types of zeros discussed by Kaul et al. (2017) because the issue of missing zeros (which is shown to be critically important in diversity analysis) was not noted in that study and zeros that appear to be outliers from empirical patterns are not considered in this study (because all random read counts are presumed to be correct).

It is typically presumed that zeros correspond to non-detected sequences, meaning that the variant is present in some quantity in the source but happened to not be included in the library and is represented by a zero. A legitimate singleton that is replaced with a zero would be a special case of a non-detect zero. Bias would result if non-detect zeros were omitted or included in the diversity analysis inappropriately (e.g., substitution with pseudo-counts or treating them as definitively absent variants). It is conceptually possible that a particular type of microorganism may be truly absent from certain sources so that the corresponding read count and proportion should definitively be zero. If false sequences due to errors in amplification and sequencing are filtered from the ASV table but left as zeros, then they are a special case of truly absent sequences. Bias would result if such zeros were included in diversity analysis in a way that manipulates them to non-zero values or allows the corresponding variant to have a plausibly non-zero proportion. Missing zeros are variants that are truly present in the source and not represented in the data—they are not acknowledged to be part of the community, even with a zero in the ASV table. Bias would result from exclusion of these zeros from diversity analysis rather than recognizing them as non-detected variants. Thus, there are three types of zeros, two of which appear indistinguishably in the data and must be handled differently and the third of which is important but does not even appear in the data. In this study, simulation-based experiments are used to illustrate implications of the dilemma of not knowing how many zeros should appear in the data to be analyzed as non-detects.

## 4.0 Probabilistic inference of source Shannon index using Bayesian methods

The Shannon index (Equation 1; Shannon, 1948; Washington, 1984) is used as a measure of alpha diversity that reflects both the richness and evenness of variants present (number of unique variants and similarity of their respective proportions). When calculated from a sample, the Shannon index (S) depends only on the proportions of the observed variants (*p*_*i*_ for the *i*^*th*^ of n variants) and not on their read counts. Critically, the Shannon index of a sample is <u>not</u> an unbiased estimate of the Shannon index of the source (even in scenarios without amplification); it is expected to increase with library size as more rare variants are observed until it converges asymptotically on the Shannon index of the source (Willis, 2019). Even if all variants in the source are reflected in the data, the precision of the estimated

Shannon index will improve with increasing library size. Building on existing work applying Bayesian methods to characterize the uncertainty in enumeration-based microbial concentration estimates (e.g., Emelko et al., 2010) and inspired by the need to consider random error in evaluation of alpha diversity that was noted by Willis (2019), a Bayesian approach is explored here for the simplified scenario of multinomially distributed data. It evaluates uncertainty in the source Shannon index given sample data, the multinomial model, and a relatively uninformative Dirichlet prior that gives equal prior weight to all variants (using a vector of ones). Hierarchical modelling that may describe how the proportional composition varies among samples is beyond the scope of this analysis. Such modelling can be beneficial when strong information in the lower tier of the hierarchy can be used to probe the fit of the upper tier; however, it can be biased if limited information in the lower tier is bolstered with flawed assumptions introduced via the upper tier.

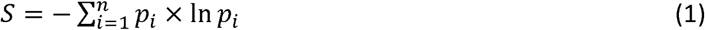

Here, a simulation study is employed that is analogous to compositional microbiome data with small library sizes and small numbers of variants and that does follow a multinomial relative abundance model. The simulation uses specified proportions for a set of variants; for illustrative purposes, the simulation represents random draws with replacement from a bag of lettered tiles based on the game Scrabble™. Randomized multinomial data (Table S1, Supplementary Content) were generated in R using varying library sizes and the proportions of the 100 tiles (including 26 letters and blanks), which correspond to a population-level Shannon index of 3.03. Markov chain Monte Carlo (MCMC) was carried out using OpenBUGS (version 3.2.3), with randomized initialization and 10,000 iterations following a 1,000-iteration burn-in. The model specification code and a small sample dataset are included in the Supplementary Content. Due to the mathematical simplicity of a multinomial model with a Dirichlet prior, this number of iterations can be completed in seconds with rapid convergence and good mixing of the Markov chain. Each iteration generates an estimate of each variant proportion, and the set of variant proportions is used to compute an estimate of the Shannon index for the source inferred from the sample data. The Markov chain of Shannon index values generated in this way is collectively representative of a sample from the posterior distribution that characterizes uncertainty in the source Shannon index given the sample data and prior. The simulated data were analyzed in several ways, as illustrated using box and whisker plots in Figure 1: (1) with all non-detected tile variants removed, (2) with zeros as needed to reach the correct number of tile variants used to simulate the data (i.e., 27), and (3) with extraneous zeros (a total of 50 tile variants of which 23 do not actually exist in the source).

**Figure 1:**
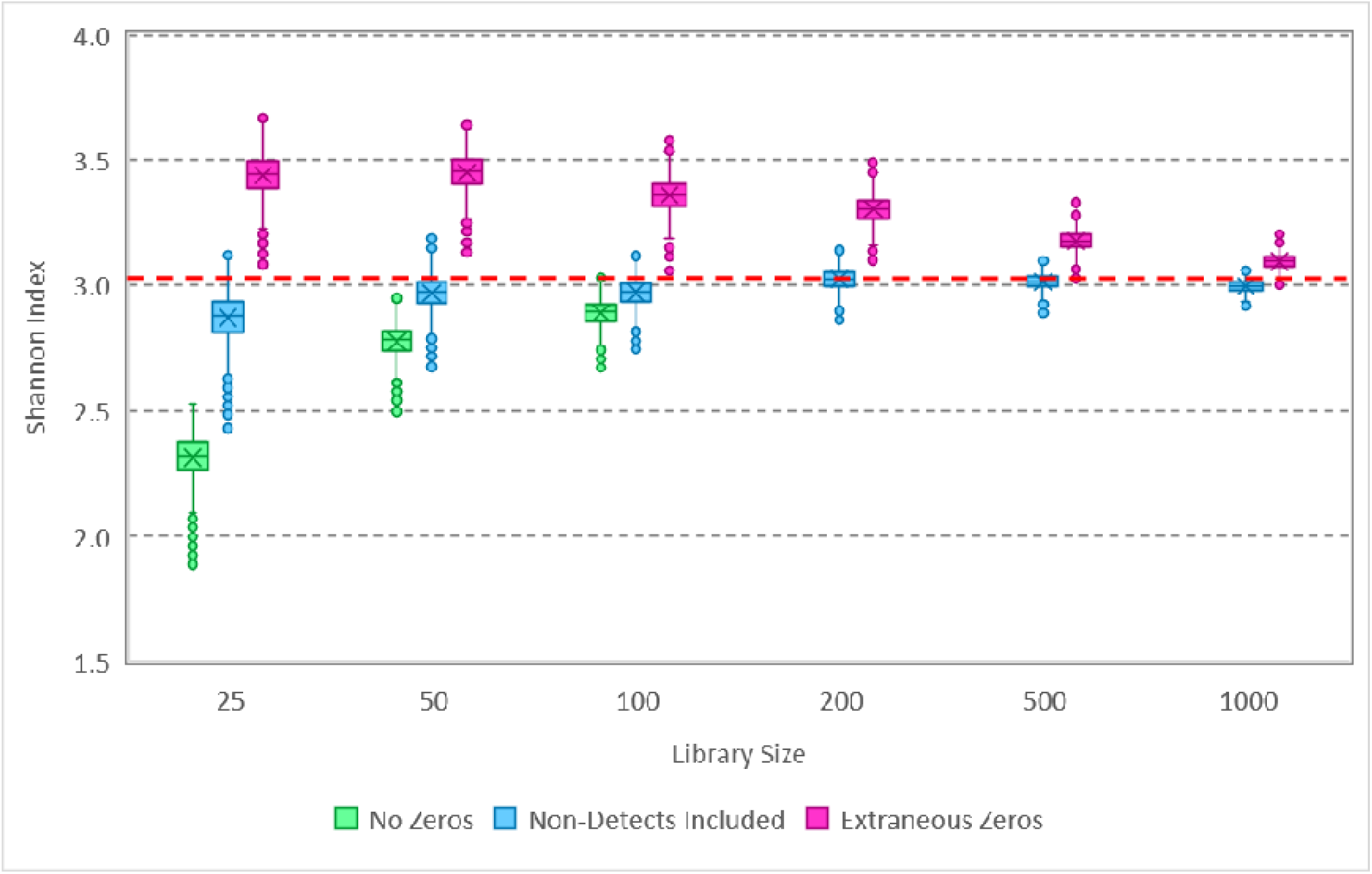
Box and whisker plot of MCMC samples from posterior distributions of the Shannon inde based on analysis of simulated data. Data with various library sizes (Table S1) were analyzed in each of 3 ways: with zeros excluded (not applicable in some cases), with zeros included for non-detected variants, and with extraneous zeros corresponding to variants that do not exist in the source. The true Shannon index of the source from which the data were simulated is 3.03.

The disparity in results between the three ways in which the data were analyzed exemplifies the importance of zeros in estimating the Shannon index of the source from which samples were gathered. Omitting non-detect zeros in this Bayesian analysis characteristically underestimates diversity, while including zeros for variants that do not exist in the source characteristically overestimates diversity. In each case, the effect diminishes as the library size is increased. Notably, the approach that included zeros for variants present in the source that were not detected in the sample allowed accurate estimation of the source Shannon index, with improving precision as the library size increases (exemplifying statistical consistency of the estimation process). Additional analysis (not shown) indicated that using a prior with a vector of 0.1’s leads to underestimation of the source Shannon inde by all three methods. Given these results, the proposed Bayesian process appears to be theoretically valid to estimate the source Shannon index from samples (for which the multinomial relative abundance model applies), and it does so without the need to normalize data with differing library sizes. Practically, however, it is not possible to know how many zeros should be included in the analysis estimating the Shannon index because the number of unique variants actually present in the source is unknown. This is a peculiar scenario that must be emphasized here because accurate statistical inference about the source is not possible: although the model form (multinomial) is known, the number of unique variants that should be included in the model is practically unknowable. Model-based supposition is not applied in this study to introduce information that is lacking; this can be a biased approach to compensating for deficiencies in observed data or flawed experiments in which “control variables” are not controlled (e.g., it is not possible to estimate concentration from a count without a measured volume) unless the supposition happens to be correct.

Because the extent to which zeros compromised accurate estimation waned with increasing library size (Figure 1), a similar analysis was performed on amplicon sequencing data for six water samples from lakes. The samples (Cameron et al., 2020) featured library sizes between 10,000 and 30,000 and observation of 1,142 unique variants among the samples. All singleton counts had been zeroed and the completed ASV table had 3,342 rows (2,200 of which are all zeros associated with variants detected in other samples from the same study area). Each sample was analyzed three ways: (1) with all non-detected sequence variants removed, (2) with zeros as needed to fill out the 1,142-row ASV table, and (3) with zeros as needed to fill out the 3,342-row ASV table. The appropriate number of zeros to be included for each sample cannot be known, but the Shannon index estimated with all non-detected sequence variants removed is very likely underestimated. The results (Figure 2) show that the number of zeros included in the analysis can have a substantial effect on the estimated Shannon index of the source, even with library sizes nearing 30,000 sequences. It is thus concluded that it is not statistically possible to estimate the Shannon index of the source (even if all the assumptions are met that enable use of the multinomial relative abundance model) unless the number of unique variants present in the source is precisely known *a priori*.

**Figure 2:**
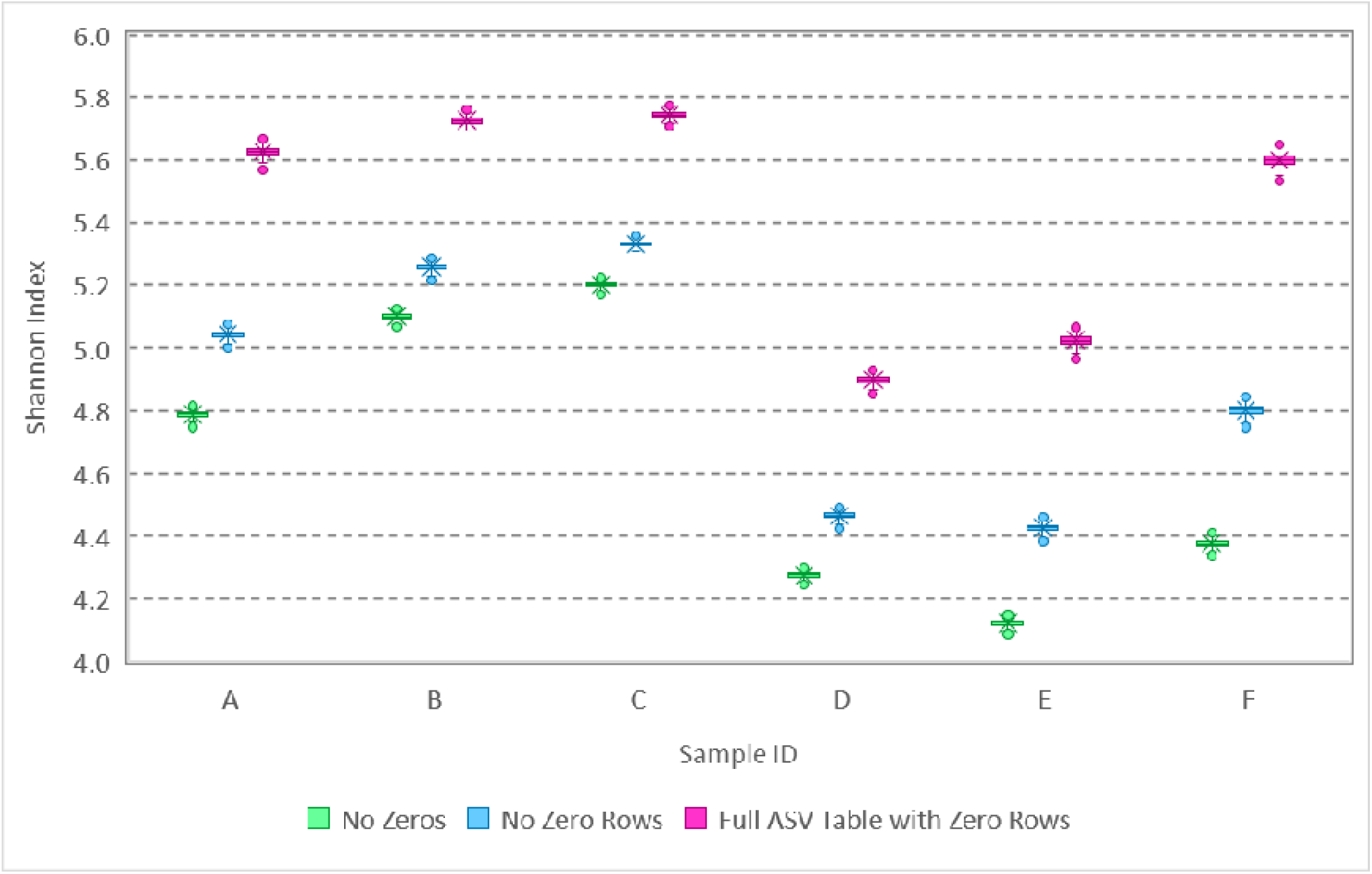
Box and whisker plot of MCMC samples from posterior distributions of the Shannon inde based on analysis of amplicon sequencing data. Data with various library sizes between 10,000 and 30,000 were analyzed 3 ways: with zeros excluded, with zeros included in a 1,142-row ASV table (no zero rows), and with additional zeros from the full 3,342-row ASV table including variants with rows of zeros (detected in other samples from the same study area).

## 5.0 Diversity Analysis in Absence of a Model to Infer Source Diversity

Recognizing that amplicon sequencing of a sample provides only partial and indirect representation of the diversity in the source (specifically partial representation of post-amplification diversity) and that statistical inference about source diversity is compromised by clustering, amplification, and not knowing how many zeros should be included in the data, the question of how to perform diversity analysis remains. The approach should recognize the random nature of amplicon sequencing data, reflect the importance of the library size in progressively revealing information about diversity, avoid normalization that distorts the proportional composition of samples, and provide some measure of uncertainty or error. Inference about source diversity is the ideal, but it is not possible with a multinomial relative abundance model unless the number of unique variants in the source is precisely known and there are many types of error in amplicon sequencing that are likely to invalidate this foundational model as discussed above. Rarefying repeatedly, a subsampling process to normalize library sizes among samples that is performed many times in order to characterize the variability introduced by rarefying (Cameron et al., 2020), satisfies these goals. When a sample is rarefied repeatedly down to a smaller library size (using sampling without replacement), it describes what data might have been obtained if only the smaller library size of sequence variants had been observed. It also does not throw out valid sequences because all sequences are represented with a sufficiently large number of repetitions. A value of the sample Shannon index may than be computed for each of the repetitions to quantify the diversity in samples of a particular library size.

Figure 3 schematically illustrates the relationship between repeatedly rarefying to smaller library sizes and statistical inference about the source from which the sample was taken. Rarefying adds random variability by subsampling without replacement while statistical inference includes parametric uncertainty that is often ignored in contemporary diversity analyses. Because the extent to which diversity is exhibited by a sample depends on the library size, such sample-level analysis must be performed at the same level (analogous to converting 1 mm, 1 cm, and 1 km to a common unit before comparing numerical values) and any observations obtained about patterns in sample-level diversity are conditional on the shared library size at which the analysis was performed. On the other hand, current methods (including rarefying once), distort the data to facilitate use of compositional analysis methods that presume the data are a perfect representation of the microbial composition in the source; it is important to recognize that the detected library is only a random sample that is imperfectly representative of source diversity.

**Figure 3:**
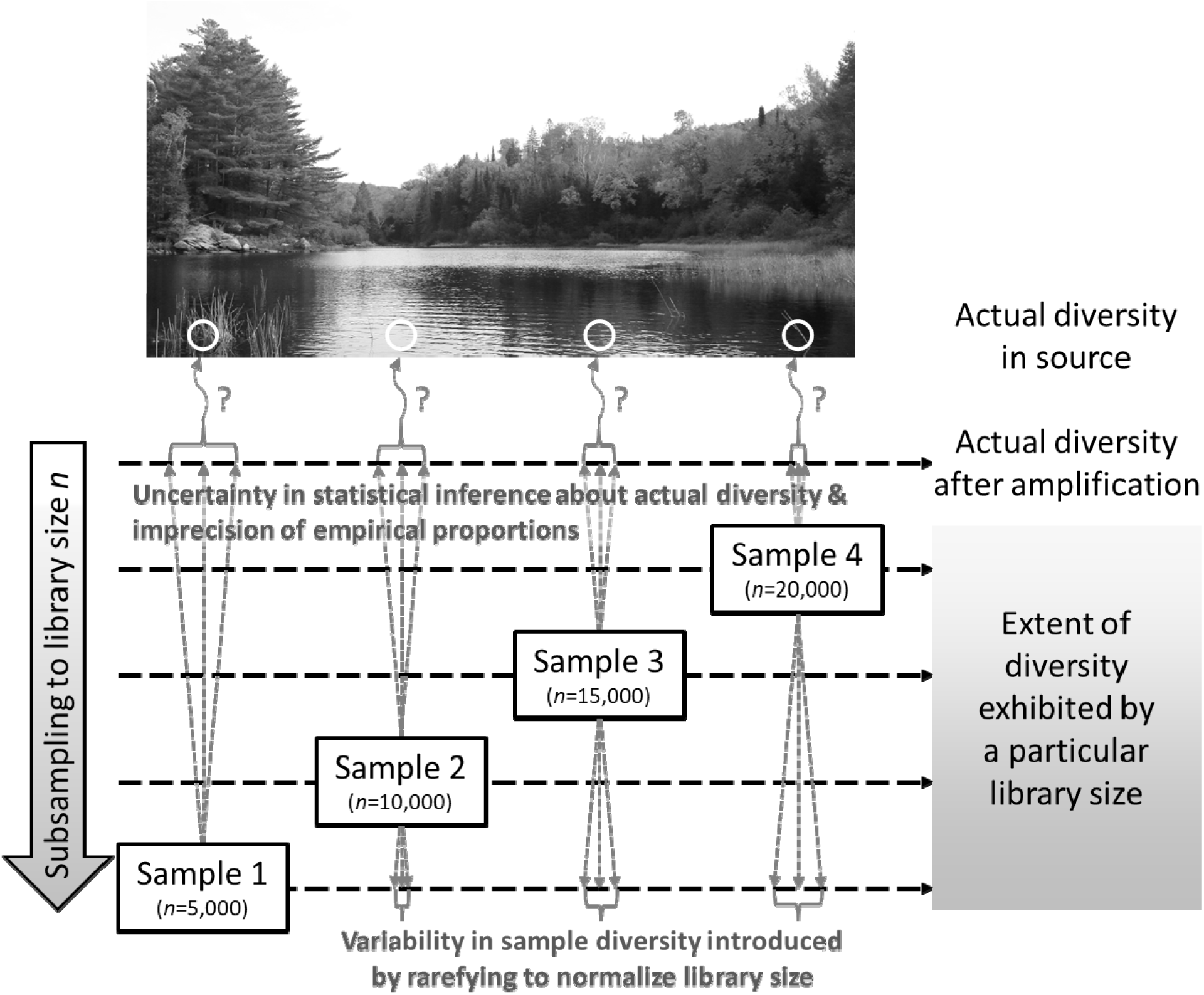
Schematic diagram relating library size and diversity quantified therefrom to uncertainty in statistical inference about source diversity and variability introduced by repeatedly rarefying to the smallest obtained library size. In this case, rarefying repeatedly evaluates the extent of the diversity (after amplification) exhibited if a library size of only *n*=5000 had been obtained from each sample.

A simulation experiment was performed using the hypothetical population based on Scrabble™ and samples with varying library sizes (see R code in Supplementary Content) to explore rarefying repeatedly and plug-in estimation of the Shannon index (Figure 4). A thousand simulated datasets with a library size of 25 yielded Shannon index values between 1.86 and 2.87 (with a mean of 2.49), illustrating that the source diversity (with a Shannon index of 3.03) is only partially exhibited by a sample with a library size of 25. Five samples were generated with library sizes of 50, 100, 200, 500, and 1000, and corresponding Shannon index values are shown in red (deteriorating markedly at library sizes of 100 or less). Each sample was then rarefied repeatedly (1000 times) to a library size of 25, resulting in the box and whisker plots of the calculated Shannon index values. Although samples with larger library sizes exhibit more diversity, samples repeatedly rarefied down to the minimum library size of 25 exhibit very comparable diversity. The Shannon index at a library size of 25 is similar for all samples, as it should be given that they were generated from the same population. If rarefying had been completed only once without quantification of the error introduced, it may erroneously have been concluded that the samples exhibited different Shannon index values.

**Figure 4:**
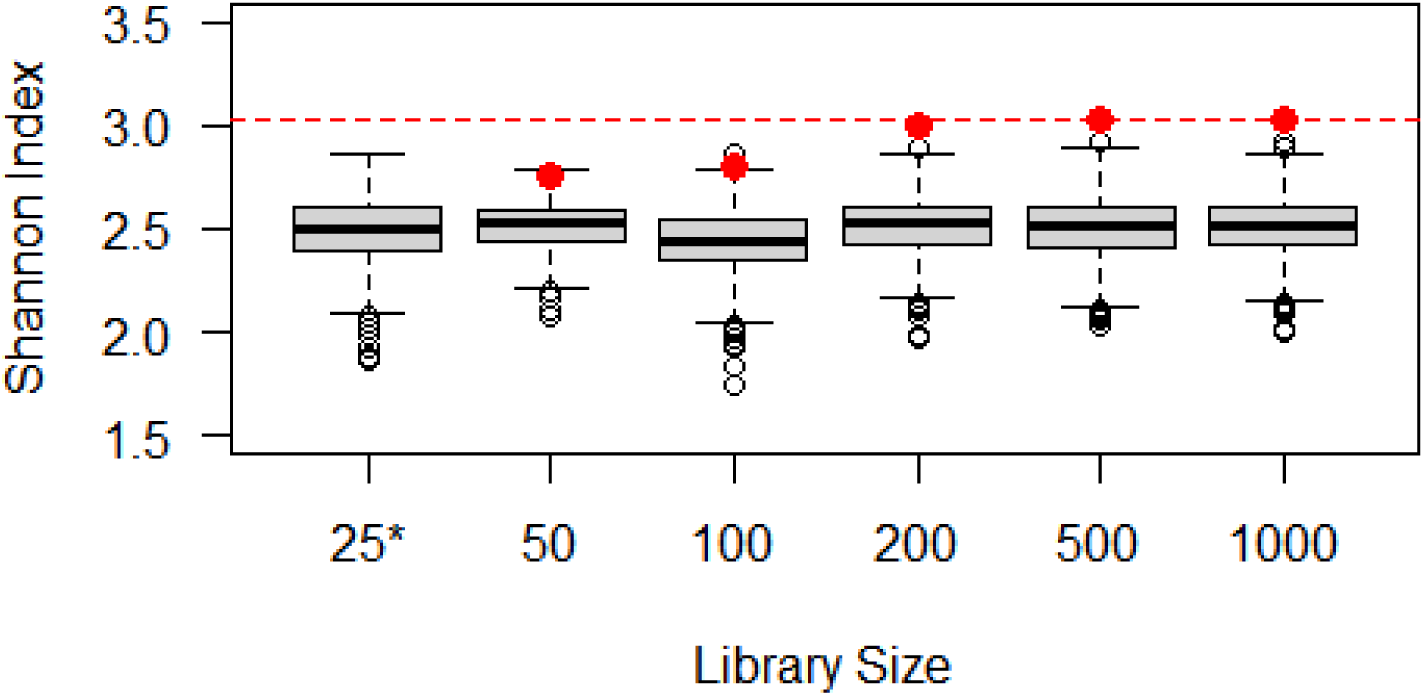
Demonstration of normalization by rarefying repeatedly using simulated data. The box and whisker plot for the library size of 25 (*) illustrates how the Shannon index varies among simulated samples and is consistently below the actual Shannon index of 3.03 (red line). The Shannon index calculated from the samples with larger library sizes (red dots) deteriorates at small library sizes. The box and whisker plots for these library sizes illustrate what Shannon index might have been calculated if only a library size of 25 had been obtained (rarefying 1000 times to this level). In all cases, a Shannon index of about 2.5 is expected with a library size of 25.

## 6.0 Discussion

Diversity analysis of amplicon sequencing data has grown rapidly, adopting tools from other disciplines but largely differing from the statistical approaches applied to classical microbiology data. Most analyses feature a deterministic set of procedures to transform the data from each sample and yield a single value of an alpha diversity metric or a single point on an ordination plot. Such procedures should not be viewed as statistical analysis because the data are not a population (i.e., perfect measurements of the proportional composition of the community in the source); they are a random sample representing only a portion of that population. Acknowledging that the data are random and that the goal is to understand the alpha and beta diversity of the sources from which samples were collected, it is important to describe and explore the error mechanisms leading to variability in the data and uncertainty in estimated diversity.

This study provides a step toward such methods by describing mechanistic random errors and their potential effects, proposing a probabilistic model and listing the assumptions that facilitate its use, discussing various types of zeros that may appear (or fail to) in ASV tables, and performing illustrative analyses using simulated data. Several sources of random error were found to invalidate the multinomial relative abundance model that is foundational to probabilistic modelling of compositional sequence count data, notably including clustering of microorganisms in the source and amplification of genes in this sequencing technology. Future simulation studies could explore the effect of non-random microorganism dispersion, sample volume (relative to a hypothetical representative elementary volume of the source), differential analytical recovery in sample processing, amplification errors, and sequencing errors on diversity analysis more thoroughly and evaluate the potential for current normalization and point-estimation approaches to misrepresent diversity.

This study also presents a simple Bayesian approach to drawing inference about diversity in the sources from which samples were collected (rather than just diversity in the sample or some transformation of it). Even under idealized circumstances in which the multinomial relative abundance model is valid, it was unfortunately found to be biased unless the number of unique variants present in the source was known a priori. This may have implications on analysis of any type of multinomial data, beyond microbiome data, in which the number of possible outcomes (or the number of outcomes with zero observations that should be included in the analysis) is unknown. It is plausible that a probabilistic model could be developed to account for errors that invalidate the multinomial model, though this would require many assumptions that would be difficult to validate and that could substantially bias inferences. In summary, probabilistic modelling should be used to draw inferences about source diversity and quantify uncertainty therein, but the simple multinomial model is invalidated by some types of error that are inherent to the method and inference is not possible even with the multinomial model unless the practically unknowable number of unique variants in the source is known.

For lack of a reliable and readily available probabilistic approach to draw inferences about source diversity, an approach to evaluate and contrast sample-level diversity at a particular library size is needed. Rarefying once manipulates the data in a way that adds variability and discards data (McMurdie & Holmes, 2014), and (like other transformations proposed to normalize data) the manipulated data are generally only used to obtain a plug-in estimate of diversity. Rarefying repeatedly, on the other hand, allows comparison of sample-level diversity estimates conditional on a library size that is common among all analyzed samples, does not discard data, and characterizes variability in what the diversity measure might have been if only the smaller library size had been observed. This approach is by no means statistically ideal, but it may be a distant second best relative to the Bayesian approach (or analogous frequentist approaches based on the likelihood function) presented in this study that cannot practically be applied in an unbiased way in many scenarios, especially due to the unknowable number of unique variants that are actually present in the source and complex error structures inherent to amplicon sequencing.

## 7.0 Acknowledgements

We acknowledge the support of NSERC (Natural Science and Engineering Research Council of Canada), specifically through the forWater NSERC Network for Forested Drinking Water Source Protection Technologies [NETGP-494312-16] and Alberta Innovates [3360-E086]. This research was undertaken, in part, thanks to funding from the Canada Research Chairs Program (M. B. Emelko; Canada Research Chair in Water Science, Technology & Policy). We are also grateful for discussion of this work with Prof. Mary E. Thompson (Department of Statistics & Actuarial Science, University of Waterloo) and assistance with performing and graphing analyses from Rachel H. M. Ruffo.

