## Supplementary Material for "Ensuring that fundamentals of quantitative microbiology are reflected in microbial diversity analyses based on next-generation sequencing"

Philip J. Schmidt^1^, Ellen S. Cameron^2^, Kirsten M. Müller^2^, Monica B. Emelko^1*^

^1^ Water Science, Technology & Policy Group, Department of Civil and Environmental Engineering, Faculty of Engineering, University of Waterloo, Waterloo, ON, Canada,

^2^ Department of Biology, Faculty of Science, University of Waterloo, Waterloo, ON, Canada,

Table S1: Simulated data based on drawing tiles from the game Scrabble^TM^ with replacement

| Tile Variant | True Proportion | Library Size | | | | | |
| --- | --- | --- | --- | --- | --- | --- | --- |
|  |  | 25 | 50 | 100 | 200 | 500 | 1000 |
| A | 0.09 | 1 | 6 | 8 | 18 | 52 | 101 |
| B | 0.02 | 0 | 1 | 2 | 4 | 7 | 12 |
| C | 0.02 | 1 | 1 | 1 | 6 | 7 | 16 |
| D | 0.04 | 1 | 0 | 5 | 6 | 20 | 40 |
| E | 0.12 | 4 | 5 | 7 | 29 | 70 | 118 |
| F | 0.02 | 0 | 0 | 2 | 6 | 13 | 22 |
| G | 0.03 | 0 | 2 | 5 | 8 | 10 | 28 |
| H | 0.02 | 1 | 2 | 1 | 3 | 5 | 20 |
| I | 0.09 | 2 | 9 | 8 | 13 | 44 | 97 |
| J | 0.01 | 0 | 0 | 2 | 2 | 1 | 11 |
| K | 0.01 | 0 | 0 | 2 | 4 | 3 | 10 |
| L | 0.04 | 1 | 2 | 1 | 3 | 28 | 52 |
| M | 0.02 | 0 | 1 | 1 | 3 | 8 | 14 |
| N | 0.06 | 0 | 5 | 7 | 10 | 27 | 51 |
| O | 0.08 | 2 | 3 | 13 | 17 | 43 | 77 |
| P | 0.02 | 1 | 1 | 2 | 6 | 9 | 18 |
| Q | 0.01 | 0 | 0 | 3 | 1 | 4 | 8 |
| R | 0.06 | 2 | 4 | 4 | 13 | 30 | 65 |
| S | 0.04 | 2 | 2 | 5 | 6 | 21 | 44 |
| T | 0.06 | 2 | 3 | 5 | 14 | 23 | 50 |
| U | 0.04 | 1 | 0 | 6 | 6 | 31 | 40 |
| V | 0.02 | 1 | 1 | 1 | 4 | 10 | 28 |
| W | 0.02 | 2 | 1 | 2 | 4 | 6 | 17 |
| X | 0.01 | 1 | 1 | 3 | 3 | 3 | 11 |
| Y | 0.01 | 0 | 0 | 1 | 4 | 8 | 19 |
| Z | 0.01 | 0 | 0 | 1 | 2 | 4 | 9 |
| Blank | 0.02 | 0 | 0 | 2 | 5 | 13 | 22 |

```
### R CODE: Simulation analysis to determine if clustering or amplification affect
### multinomial model for amplicon sequencing
```

```
### Install package ggplot2 and add to library
install.packages("ggplot2")
library(ggplot2)

### Fix randomization seed for reproducibility
set.seed(20210528)

### ANALYSIS #1 – CLUSTERING

### Generate dataset (10^6 draws) from A0 ~ binomial(n=6, p=2/3) & B0 = 6 – A0
### Special case of multinomial relative abundance model in which A:B = 2:1
A0 <- rbinom(1000000, 6, 2/3)
B0 <- 6 - A0

### Construct a negative binomial model as a function of the mean (mu=rho*lambda)
### and over-dispersion relative to Poisson (lambda) in which A:B = 2:1
### Simple model with microorganisms clustering within but not between varieties
### Generate dataset (10^6 draws) from A1 ~ negbin(size=40,000, prob=10,000/10,001)
### & B1 ~ negbin(size=20,000, prob=10,000/10,001)
### With lambda = 0.0001, variants are essentially not clustered and the negative
### binomial model is approximately Poisson(mu)
```

```
muA <- 4
lambdaA <- 0.0001
rhoA <- muA / lambdaA
sizeA <- rhoA
probA <- 1/(1 + lambdaA)
muB <- 2
lambdaB <- 0.0001
rhoB <- muB / lambdaB
sizeB <- rhoB
probB <- 1/(1 + lambdaB)
A1 <- rnbinom(1000000, size = sizeA, prob = probB)
B1 <- rnbinom(1000000, size = sizeB, prob = probB)
SUM1 <- A1 + B1
### Create data frame of subset of simulations in which SUM1 = 6
Poisson <- subset(data.frame(A1, B1, SUM1), SUM1 == 6)
```

```
### Generate dataset (10^6 draws) from A2 ~ negbin(size=4/3, prob=0.25)
### & B2 ~ negbin(size=2/3, prob=0.25)
### With lambda = 3, variants are substantially clustered
muA <- 4
lambdaA <- 3
rhoA <- muA / lambdaA
sizeA <- rhoA
probA <- 1/(1 + lambdaA)
muB <- 2
lambdaB <- 3
rhoB <- muB / lambdaB
sizeB <- rhoB
probB <- 1/(1 + lambdaB)
A2 <- rnbinom(1000000, size = sizeA, prob = probB)
B2 <- rnbinom(1000000, size = sizeB, prob = probB)
SUM2 <- A2 + B2
### Create data frame of subset of simulations in which SUM2 = 6
Cluster <- subset(data.frame(A2, B2, SUM2), SUM2 == 6)


### Construct graph of relative frequency of A0, A1, and A2
A0relfreq <- table(A0)/1000000
A1relfreq <- table(Poisson$A1)/length(Poisson$A1)
A2relfreq <- table(Cluster$A2)/length(Cluster$A2)
Legend <- rep(c("A0", "A1", "A2"), each = 7)
X <- rep(c(0, 1, 2, 3, 4, 5, 6), times = 3)
Y <- c(A0relfreq, A1relfreq, A2relfreq)
summary1 <- data.frame(Y, Legend, X)
ggplot(summary1, aes(fill = Legend, y = Y, x = X)) +
  labs(y = "Relative Frequency", x = "Count A") +
  scale_fill_manual(values = c("#66E199", "#66CCFF", "#FF33CC")) +
  geom_bar(position = "dodge", stat = "identity")
```

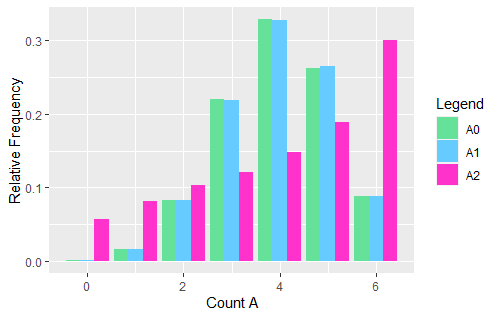

```
### Relative frequencies of A0 and A1 are similar, suggesting that random sampling ### error following a Poisson model is compatible with the multinomial relative
### abundance model. Relative frequencies of A0 and A2 are dissimilar, proving that ### random sampling error following a negative binomial model is incompatible with
### the multinomial relative abundance model.


### ANALYSIS #2 - AMPLIFICATION

### Generate dataset (10^6 draws) from A3 ~ binomial(n=4, p=2/3) & B3 = 4 – A3
### Special case of multinomial relative abundance model in which A:B = 2:1
A3 <- rbinom(1000000, 4, 2/3)
B3 <- 4 - A3
### Amplify A3 and B3 with a 50% success rate
A3Amp <- A3 + rbinom(1000000, A3, 0.5)
B3Amp <- B3 + rbinom(1000000, B3, 0.5)
SUM3 <- A3Amp + B3Amp
### Create data frame of subset of simulations in which SUM3 = 6
Amplify <- subset(data.frame(A3Amp, B3Amp, SUM3), SUM3 == 6)

### Construct graph of relative frequency of A0 and A3
A3relfreq <- table(Amplify$A3Amp)/length(Amplify$A3Amp)
Legend <- rep(c("A0", "A3"), each = 7)
X <- rep(c(0, 1, 2, 3, 4, 5, 6), times = 2)
Y <- c(A0relfreq, A3relfreq)
summary2 <- data.frame(Y, Legend, X)
ggplot(summary2, aes(fill = Legend, y = Y, x = X)) +
  labs(y = "Relative Frequency", x = "Count A") +
  scale_fill_manual(values = c("#66E199", "#66CCFF")) +
  geom_bar(position = "dodge", stat = "identity")
```

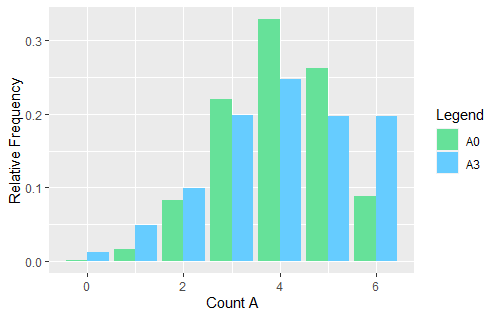

```
### Relative frequencies of A0 and A3 are dissimilar, proving that amplification is
### incompatible with the multinomial relative abundance model.

OpenBUGS Code for Multinomial Relative Abundance Data

p[i]: fraction of sequences of type i - STOCHASTIC (parameter of interest)
X[i]: count of sequences of type i - STOCHASTIC (measured data)
PlnP[i]: p*ln(p) of type i - DETERMINISTIC
Shannon: Shannon index evaluated using posterior proportions - DETERMINISTIC

model {
	# Define prior that is equal for all fractions in the multinomial model
	p[1:m] ~ ddirich(alpha[])
	# Define likelihood
	X[1:m] ~ dmulti(p[1:m], N)
	N <- sum(X[])
	for (i in 1:m) {
		PlnP[i] <- p[i]*log(p[i])
	}
	Shannon <- -sum(PlnP[])
}

DATA - Simple example with four variants
list(m=4)
alpha[]	X[]
1	40
1	30
1	20
1	10
END

### R CODE: Generation of Figure 4 (sample-level Shannon index simulation analysis)

### Generate population (based on Scrabble)
Variants <- c("A", "B", "C", "D", "E", "F", "G", "H", "I", "J", "K", "L", "M",
  "N", "O", "P", "Q", "R", "S", "T", "U", "V", "W", "X", "Y", "Z", "BLANK")
Fractions <- c(0.09, 0.02, 0.02, 0.04, 0.12, 0.02, 0.03, 0.02, 0.09, 0.01, 0.01,
  0.04, 0.02, 0.06, 0.08, 0.02, 0.01, 0.06, 0.04, 0.06, 0.04, 0.02, 0.02, 0.01,
  0.02, 0.01, 0.02)
ShannonPop <- -sum(Fractions * log(Fractions))

### Generate 1000 samples with a library size of 25, compute Shannon Index
set.seed(44335)   # Fixed seed for randomizations
Data25 <- rmultinom(n = 1000, size = 25, prob = Fractions)
Shannon25 <- -colSums(Data25 / 25 * log(Data25 / 25), na.rm = TRUE)

### Generate 5 samples with larger library sizes, compute Shannon index
Data50 <- rmultinom(n = 1, size = 50, prob = Fractions)
Data100 <- rmultinom(n = 1, size = 100, prob = Fractions)
Data200 <- rmultinom(n = 1, size = 200, prob = Fractions)
Data500 <- rmultinom(n = 1, size = 500, prob = Fractions)
Data1000 <- rmultinom(n = 1, size = 1000, prob = Fractions)
ShannonRaw50 <- -sum(Data50 / 50 * log(Data50 / 50), na.rm = TRUE)
ShannonRaw100 <- -sum(Data100 / 100 * log(Data100 / 100), na.rm = TRUE)
ShannonRaw200 <- -sum(Data200 / 200 * log(Data200 / 200), na.rm = TRUE)
ShannonRaw500 <- -sum(Data500 / 500 * log(Data500 / 500), na.rm = TRUE)
ShannonRaw1000 <- -sum(Data1000 / 1000 * log(Data1000 / 1000), na.rm = TRUE)

### Repeatedly rarefy data with large library sizes to rarefied library size 25
library(extraDistr)
Rarefy50 <- rmvhyper(nn = 1000, n = t(Data50), k = 25)
Shannon50 <- -rowSums(Rarefy50 / 25 * log(Rarefy50 / 25), na.rm = TRUE)
Rarefy100 <- rmvhyper(nn = 1000, n = t(Data100), k = 25)
Shannon100 <- -rowSums(Rarefy100 / 25 * log(Rarefy100 / 25), na.rm = TRUE)
Rarefy200 <- rmvhyper(nn = 1000, n = t(Data200), k = 25)
Shannon200 <- -rowSums(Rarefy200 / 25 * log(Rarefy200 / 25), na.rm = TRUE)
Rarefy500 <- rmvhyper(nn = 1000, n = t(Data500), k = 25)
Shannon500 <- -rowSums(Rarefy500 / 25 * log(Rarefy500 / 25), na.rm = TRUE)
Rarefy1000 <- rmvhyper(nn = 1000, n = t(Data1000), k = 25)
Shannon1000 <- -rowSums(Rarefy1000 / 25 * log(Rarefy1000 / 25), na.rm = TRUE)

### Plotting
boxplot(Shannon25, Shannon50, Shannon100, Shannon200, Shannon500, Shannon1000,
  xlab = "Library Size", ylab = "Shannon Index", ylim = c(1.5,3.5), las = 1,
  names = c("25*", "50", "100", "200", "500", "1000"))
### Add sample Shannon index values
  abline(ShannonPop, 0, col = "red", lty = 2)
  points(2, ShannonRaw50, col = "red", pch = 16, cex = 1.25)
  points(3, ShannonRaw100, col = "red", pch = 16, cex = 1.25)
  points(4, ShannonRaw200, col = "red", pch = 16, cex = 1.25)
  points(5, ShannonRaw500, col = "red", pch = 16, cex = 1.25)
  points(6, ShannonRaw1000, col = "red", pch = 16, cex = 1.25)
```
